# HIV-1 Gag protein with or without p6 specifically dimerizes on the viral RNA packaging signal

**DOI:** 10.1101/2020.06.18.159822

**Authors:** Samantha Sarni, Banhi Biswas, Shuohui Liu, Erik D. Olson, Jonathan P. Kitzrow, Alan Rein, Vicki H. Wysocki, Karin Musier-Forsyth

## Abstract

The HIV-1 Gag protein is responsible for genomic RNA (gRNA) packaging and immature viral particle assembly. While the presence of gRNA in virions is required for viral infectivity, in its absence, Gag can assemble around cellular RNAs and form particles resembling gRNA-containing particles. When gRNA is expressed, it is selectively packaged despite the presence of excess host RNA, but how it is selectively packaged is not understood. Specific recognition of a gRNA packaging signal (Psi) has been proposed to stimulate the efficient nucleation of viral assembly. However, the heterogeneity of Gag-RNA interactions renders capturing this transient nucleation complex using traditional structural biology approaches challenging. Here, we used native mass spectrometry to investigate RNA binding of wild-type Gag and Gag lacking the p6 domain (GagΔp6). Both proteins bind to Psi RNA primarily as dimers, but to a control RNA primarily as monomers. The dimeric complexes on Psi RNA require an intact dimer interface within Gag. GagΔp6 binds to Psi RNA with high specificity *in vitro* and also selectively packages gRNA in particles produced in mammalian cells. These studies provide direct support for the idea that Gag binding to Psi specifically nucleates Gag-Gag interactions at the early stages of immature viral particle assembly in a p6-independent manner.

## Introduction

The ability to specifically select the viral genomic RNA (gRNA) for packaging into the assembling virus particle is absolutely necessary for specific replication of HIV-1 or other retroviruses. This selection is critical as the gRNA is surrounded by a great excess of cellular RNAs, and these RNAs can also be packaged under certain conditions. Approximately 2500 copies of the viral structural protein (“Gag”) assemble around a gRNA dimer forming an immature virion (1–5). The full-length 9.4 kb viral RNA that is packaged into virions serves both as as gRNA and as the mRNA for translation of all structural and enzymatic viral proteins including Gag (6). Remarkably, when expressed in cells lacking viral gRNA, Gag still forms virus-like particles (VLPs) (7). Thus, despite the critical role of gRNA for the infectivity of viral particles, Gag VLP assembly does not depend on its presence. While there are many therapeutics in clinical use that target various steps of the viral lifecycle including entry, reverse transcription, and integration, there are currently no antiviral therapies that inhibit gRNA packaging or virion assembly (8).

The mechanism of selective packaging of HIV-1 gRNA is not well understood. The selection depends upon its “packaging signal” (Psi), a region of ∼ 100 bases near the 5′ end of the RNA. We have recently found that at physiological ionic strengths *in vitro*, Gag binds with roughly equal affinity to Psi-containing and control RNAs (9, 10). In light of these observations, we and others (7, 11, 12) have suggested that gRNA is selectively packaged because it initiates assembly more rapidly or more efficiently than other RNAs. Understanding the initial stages of this process, i.e., nucleation of Gag-Gag interactions, is an essential step in developing new therapeutics that can interfere with immature particle assembly.

One complication in the attempt to reconcile the selective packaging observed *in vivo* with the *in vitro* binding data is that the recombinant Gag protein used *in vitro* has, with very few exceptions (13–16), lacked the C-terminal “p6” domain. The Gag protein is composed of four major functional domains from N-to C-terminus: matrix (MA), capsid (CA), nucleocapsid (NC), and p6, as well as the short spacer peptides SP1 and SP2 (Figure 1A). MA is cotranslationally myristoylated (17), and during virion assembly this domain becomes anchored to the plasma membrane via its myristoyl group (18, 19). MA is also highly basic and capable of binding RNA (18, 20–25). Gag-Gag interactions are mediated by CA and SP1, and the NC domain is primarily responsible for specific gRNA recognition (26, 27). The p6 domain facilitates the release of viral particles from the surface of virus-producing cells, and is not, as far as is known, involved in the formation of the virus (2); it has frequently been omitted from the Gag protein produced in bacteria largely for reasons of convenience. However, it has recently been suggested that p6 may, in fact, function in the selective packaging of gRNA or Gag-Gag oligomerization during viral assembly (14, 15). The first goal of the experiments presented here is to determine whether the p6 domain is dispensable in the selective packaging of gRNA by Gag.

**Figure 1.**
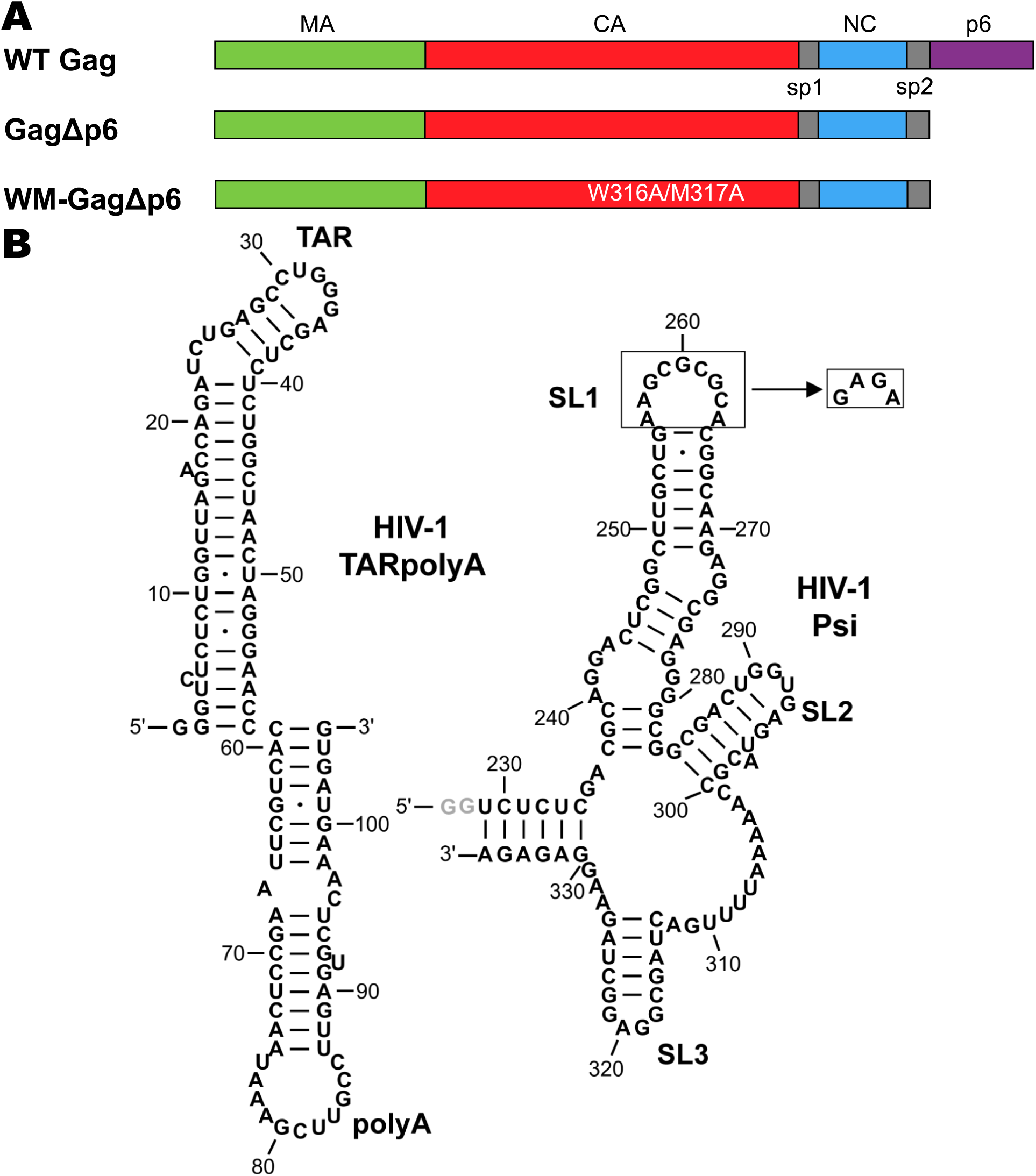
(A) Domain structure of WT and mutant Gag proteins investigated in this work. (B) Sequence and predicted secondary structures of TARpolyA (*left*) and Psi (*right*) RNAs. To ensure a monomeric state, the SL1 loop of Psi was mutated to a GAGA tetraloop as indicated. The gray nucleotides at the 5’ end of Psi (GG) are not encoded by HIV-1 but were added to improve the yield of *in vitro* transcription.

In the cytosol, HIV-1 Gag exists as monomers or lower-order oligomers and only forms higher order multimers at the plasma membrane (28–30). Packaging of gRNA is initiated in the cytoplasm through specific, non-electrostatic interactions between the NC domain of Gag and Psi (7, 22, 31, 32). The 5′UTR regulates many stages of the viral lifecycle including genome dimerization, splicing, and initiation of reverse transcription, as well as packaging. In addition to Psi, the 5′UTR is composed of several structural elements: the transactivation response hairpin (TAR), the poly-A hairpin (polyA), and the primer binding site (PBS). Psi is composed of three stemloops (SL1, SL2, SL3) with a conserved GC-rich palindromic dimerization initiation site (DIS) located within SL1. Although dimerization of gRNA appears to be required for its packaging *in vivo* (12, 33), whether specific interaction of Gag with Psi depends on dimerization *in vitro* is less clear (7, 9, 10, 16, 34–36). Because Gag can assemble around non-gRNA in the cell, we hypothesize that Gag-Gag interactions are facilitated by NC domain binding to Psi in a manner that is distinct from binding to non-Psi RNA sequences. In this work, native mass spectrometry (nMS) was carried out to test this hypothesis. We used a 109-nt Psi construct that was previously characterized to display high Gag binding specificity, and the transactivation response-polyA hairpin element (TARPolyA or TpA) as the non-Psi seqeunce, as this RNA was previously shown to be characterized by low Gag binding specificity (10). The second major goal of this work is to investigate the role of Psi (vs. TpA) RNA binding in nucleating higher order Gag multimerization. The dimerization of Gag is mediated by an interface within CA. To determine whether complexes comprised of two Gag and one RNA molecule were formed via Gag-Gag as well as Gag-RNA interactions, we tested a Gag mutant (“WM-GagΔp6”) lacking this interface, using *in vitro* binding assays, nMS, and cell-based RNA packaging assays. The results strongly support the hypothesis that Gag-Gag interactions are facilitated by binding to Psi and that these phenomena are all independent of the p6 domain.

## Results

### Gag oligomeric state in the absence of RNA

nMS was first used to analyze the purity of the Gag constructs studied in this work (Figure 1A). The experimental molecular masses for the monomeric and dimeric forms of each Gag construct closely match the theoretical molecular masses, confirming the high purity of the preparations and the binding of two zinc ions (Figure S1-S3,Table S1). We next examined the oligomeric state of free Gag proteins at multiple concentrations in the absence of any nucleic acids. Wild-type (WT) Gag and GagΔp6 were largely monomeric under these conditions, but contained a small fraction of dimers at concentrations > 6 µM (Figure 2 panels A1, B1; Figure S1 and S2). While the presence of the p6 domain had minimal effect on the oligomeric states observed, it did influence the charge state distribution. The multimodal charge state distribution observed for WT Gag is attributed to the presence of the intrinsically disordered p6 domain (37). As expected, dimerization depended on the well-characterized dimer interface in the C-terminal domain of CA, as WM-GagΔp6 was exclusively monomeric (Figures 2C1 and S3) (38, 39).

**Figure 2.**
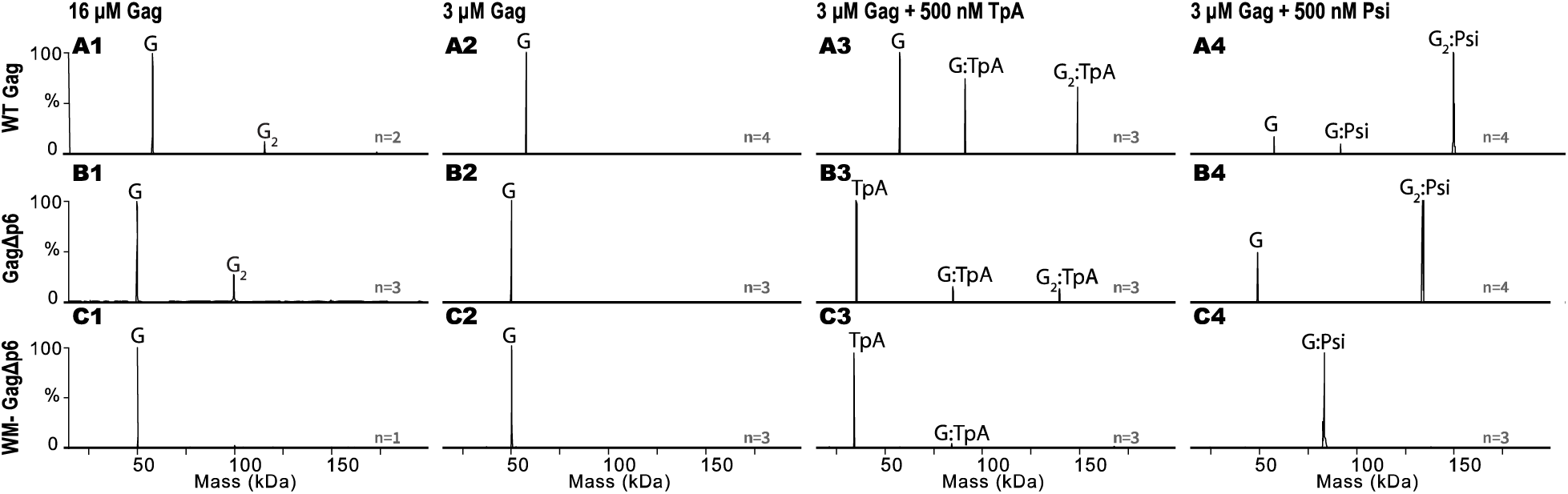
Zero-charge mass spectra of: WT Gag (A1), GagΔp6 (B1) and WM-GagΔp6 (C1) at 16 µM; WT Gag (A2), GagΔp6 (B2) and WM-GagΔp6 (C2) at 3 µM; 500 nM TARpolyA in the presence of 3 µM WT Gag (A3), GagΔp6 (B3) and WM-GagΔp6 (C3); 500 nM Psi in the presence of 3 µM WT Gag (A4), GagΔp6 (B4) and WM-GagΔp6 (C4). Notation is as follows: G=Gag, G_2_=two Gag, TpA=TARpolyA. Molecular masses for all analytes are listed in Table S1. The number of replicates (n) per experiment are indicated on each spectrum with additional repeats shown in Supplemental Figures S1-S6.

### Gag oligomeric state in the presence of RNA

We also analyzed the oligomeric state of Gag in the presence of Psi RNA and a control RNA, TARpolyA (Figure 1B). Both WT Gag and GagΔp6 formed complexes with TARpolyA. These complexes were found to contain either one or two copies of Gag and a single copy of TARpolyA (Figure 2, panels A3, B3, Figures S4, and S5). In contrast, in the presence of Psi these proteins formed complexes comprised almost entirely of two copies of Gag bound to one copy of Psi RNA (Figure 2, panels A4, B4; Figures S4 and S5). The presence of the p6 domain had no significant effect on the protein’s oligomeric state in the presence of either RNA, although the amount of the complexes detected in the presence of TARpolyA was greater for the full-length protein (Figure 2 row A vs. row B). The relative abundance of the observed species varied between replicates, but the observed differences in Gag:Psi relative to Gag:TARpolyA binding stoichiometry remained consistent (Figures S4-S6).

The formation of Gag:RNA complexes with 2:1 stoichiometry may be explained either by Gag binding to RNA as a dimer, or alternatively by the two Gag molecules binding to two separate RNA binding sites. To distinguish between these two possibilities, we used WM-GagΔp6. WM-GagΔp6 formed complexes containing one copy of WM-GagΔp6 and one copy of RNA for both TARpolyA and Psi RNAs (Figure 2, panels C3, C4, and Figure S6), supporting the conclusion that Psi RNA binding promotes dimerization of Gag rather than cooperative Gag binding to two independent binding sites in Psi RNA. More WM-GagΔp6 protein was bound to Psi than to TARpolyA RNA. Collectively, these observations demonstrate that the primarily monomeric WT Gag protein dimerizes by protein-protein interaction, using the “WM” interface, in the presence of RNA. We propose that the increased abundance of Gag dimer in the presence of Psi RNA results from changes in Gag conformation upon NC domain binding to Psi, which exposes dimerization interfaces. Previous studies have demonstrated that the mature NC domain can adopt different conformations upon binding to different RNAs (40, 41). Thus, it is feasible that the identity of the RNA dictates the conformation NC adopts when bound, which may allosterically regulate the formation of a dimerization competent Gag conformation.

### Similar Psi RNA binding specificity for GagΔp6 and WT Gag *in vitro*

To further investigate any differences in binding specificity between WT Gag and GagΔp6, we performed fluorescence anisotropy (FA)-based salt-titration binding assays (42). A non-specific interaction between protein and RNA, mediated primarily by electrostatic interactions, will dissociate at a lower salt concentration than one that also involves specific non-electrostatic interactions (Figure 3). Two parameters can be determined from this analysis: K_d,1M_ and Z_eff_. The extrapolated dissociation constant at 1 M salt (K_d,1M_) reflects the strength of non-electrostatic contributions to binding, while Z_eff_ is a measure of the number of Na^+^ ions displaced from RNA upon protein binding (43, 44). Using this assay, we previously reported that the binding of GagΔp6 to Psi is far more salt-resistant than its binding to TARpolyA (10). In good agreement with these previous studies, the K_d,1M_ values for GagΔp6 binding to Psi and TARpolyA RNAs are 3.9 × 10^−5^ M and 1.5 × 10^−1^ M, respectively, and the values of Z_eff_ are 6.1 and 10.8, respectively. In the case of WT Gag, the K_d,1M_ values for Psi and TARpolyA are 7.7 × 10^−5^ M and 3.4 × 10^−1^ M, respectively, and the Z_eff_ values are 6.3 and 10.4, respectively. Thus, we conclude that the presence of the p6 domain does not significantly affect the specificity of Gag towards Psi RNA (Figure 3). Importantly, titrations using ammonium acetate in place of the typical NaCl were also performed, which verified that significant binding differences between Psi and TARPolyA RNA are observed under nMS-compatible solution conditions (Figure S7 and S8).

**Figure 3.**
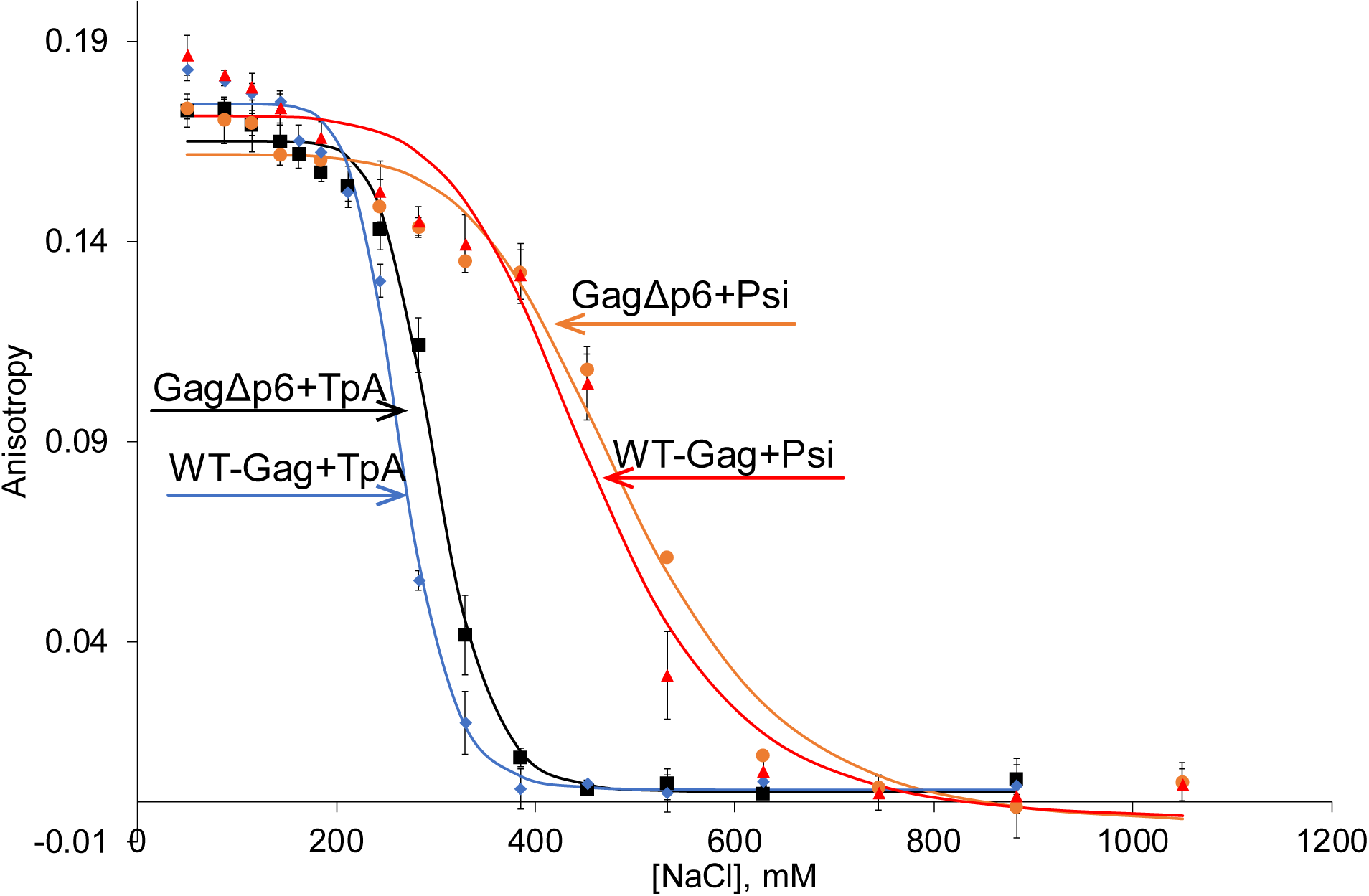
Fluorescence anisotropy results for WT Gag and GagΔp6 binding to Psi and TARPolyA RNAs as a function of NaCl concentration (n=3, error bars reflect the standard deviation of three independent measurements).

### Selective packaging of gRNA in cells is independent of p6

We also tested the ability of Gag with and without the p6 domain to selectively package gRNA in human cells. 293T cells were transiently transfected with our Gag expression plasmid and a plasmid expressing an HIV-derived GFP vector. This vector contains Psi and will therefore be selectively packaged by WT Gag. Packaging of the vector RNA in released virus particles was quantitated by real-time RT-PCR. Parallel experiments were performed using a mutant Gag expression plasmid in which the Gag coding region lacked the p6 domain.

The effects of p6 removal upon virus particle production were quantified by Western blot analysis of viral pellets from the medium of the transfected cultures. GagΔp6 produced a lower level of VLPs than WT Gag. The average yield of VLPs over several experiments with GagΔp6 was 45% of that obtained in parallel transfections of WT Gag. Similar levels of WT Gag and GagΔp6 were present in lysates of the transfected cells (Figure 4), showing that the expression of the two proteins was similar. Transmission electron micrographs of cells expressing GagΔp6 (Figure S9) showed many partially formed and malformed VLPs, as well as assembled particles arrested at the cell surface. All of these observations are fully consistent with the known function of p6 in virus particle release (27, 45, 46).

**Figure 4.**
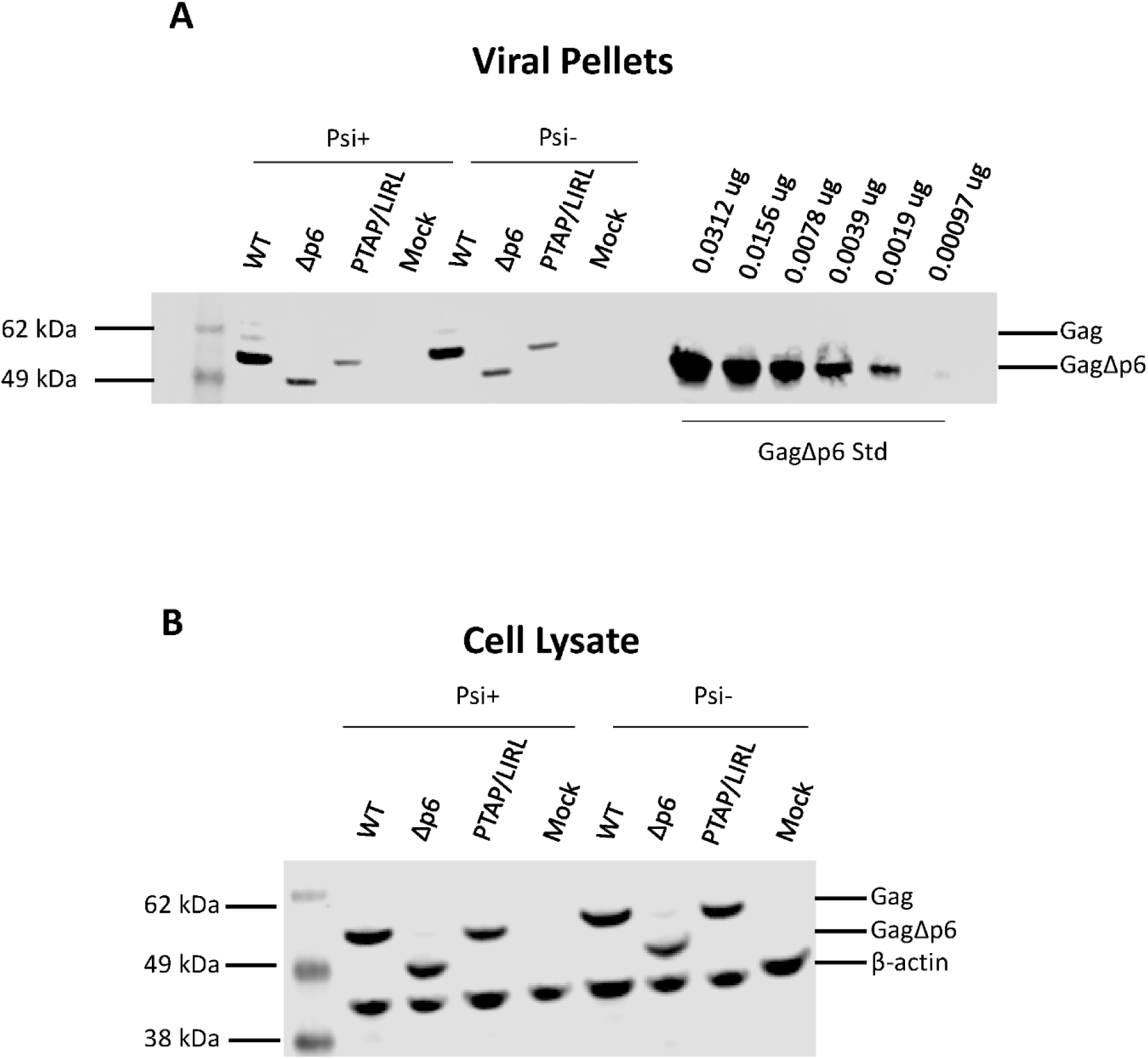
Viral pellets (A) and cell lysates (B) from a representative transfection experiment were analyzed for Gag by Western blot analysis as described in the Experimental Procedures. The cell lysates (B) were also probed for β-actin. A dilution series of recombinant GagΔp6 protein was loaded adjacent to the viral pellets (A) so that a standard curve could be constructed (Figure S12).

RNA was also extracted from the pellets and assayed for the vector RNA; the results of these measurements, corrected for the difference in virus particle content, are shown in Figure 5A. It is evident that the ratio of vector RNA to Gag protein in the viral pellets from GagΔp6 is only ∼ 4% of the ratio in the WT Gag pellets.

**Figure 5.**
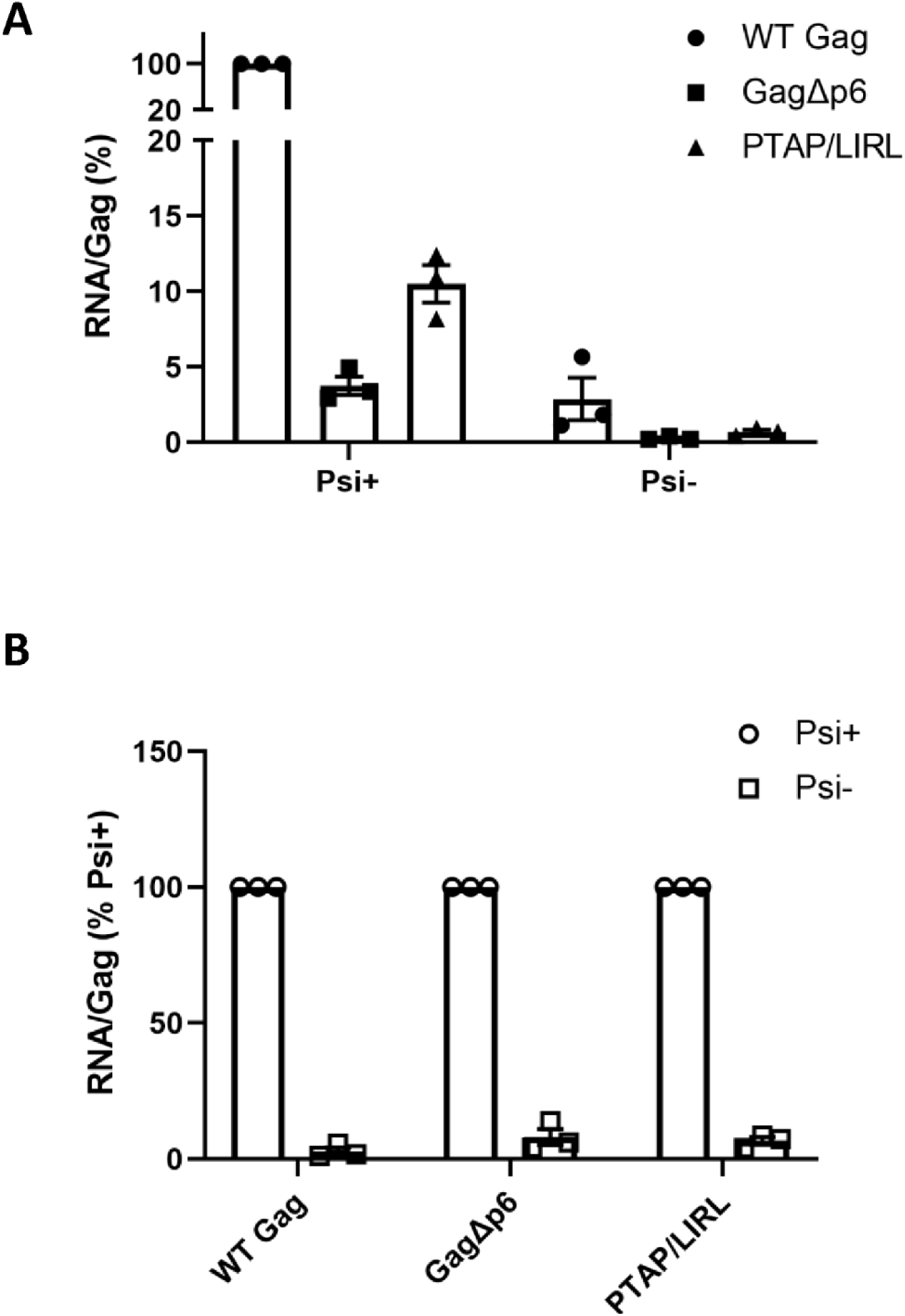
RNA packaging by mutant Gag proteins. The figure shows the results of three separate transfections. In each case, viral pellets were assayed for copies of the vector RNA and for Gag content as described in the Experimental Procedures. A: The ratio of these two quantities in the WT control was set to 100. B: For each Gag, the ratio of these two quantities for Psi^+^ was set to 100.

The low level of vector RNA in the GagΔp6 pellets might indicate that GagΔp6 has completely lost the ability to selectively package Psi-containing RNA; perhaps this level represents non-specific packaging, as seen with nearly any cellular mRNA in the absence of Psi-specific packaging of gRNA (47). Alternatively, the ability to selectively package Psi-containing RNA might be retained but somewhat diminished in GagΔp6. To distinguish between these possibilities, we measured the packaging of a derivative of the vector lacking Psi (Psi^-^). As shown in Figure 5B, the deletion of Psi profoundly reduced the packaging of the vector RNA by GagΔp6 (as well as by WT Gag), so that the RNA:Gag ratio in the pellets was < 10% of that seen with the Psi+ vector. These data show that the GagΔp6 still retains, to a very significant extent, the ability to preferentially package Psi-containing RNA.

The p6 domain functions in the interactions of Gag with the cellular ESCRT machinery during virus particle budding from virus-producing cells. Thus, it is conceivable that the ablation of this interaction in GagΔp6 is somehow responsible for the reduction in packaging of Psi^+^ RNA. Indeed, as noted above, malformed or partial virions can be seen in electron micrographs of transfected cells (Figure S9); perhaps some of these are released without packaging gRNA, or perhaps the malformed GagΔp6 particles fail to protect their RNA cargo from degradation. To explore this possibility, we also measured packaging of the vector RNA by full-length Gag in which the PTAP motif, the site in p6 that interacts with the ESCRT component Tsg101, was replaced with LIRL (46). This mutation, like the removal of p6, modestly reduced the level of particle production (Figure 4) and resulted in the formation of misshapen particles (Figure S9). As shown in Figure 5A, these pellets also show a far lower ratio of vector RNA to Gag protein than those formed by WT Gag. As with WT and GagΔp6, we also found that removal of Psi from the vector significantly reduced its packaging by the PTAP mutant Gag, showing that the packaging of the intact vector by this mutant Gag is Psi-specific (Figure 5B). Taken together, the data suggest that the reduction in Psi-specific RNA packaging by GagΔp6 is a result of its defective interaction with ESCRT machinery, and that it has not lost specificity in its interactions with RNA. It has previously been reported that virions assembled from GagΔp6 or PTAP-Gag are deficient in reverse transcriptase and integrase (46); it seems likely that gRNA, like these internal viral proteins, is lost from or degraded within the budding mutant virions during the extended period between assembly and release from the cell.

## Conclusions

The data presented here lead to two main conclusions regarding the interactions between Gag, the structural protein of HIV-1 virus particles, and RNA. First, the presence or absence of the p6 domain has minimal effect on Psi interactions in *in vitro* binding assays, and does not appear to significantly affect the packaging of viral RNA in virus-producing cells, except for an effect attributable to the “late domain” mutant phenotype seen with this truncated Gag protein. Second, Gag (with or without p6) dimerization is promoted when it binds the Psi RNA packaging signal, but far less dimer is observed when it binds a control RNA (Figure 2).

A significant effect of the p6 domain on Gag/RNA interactions, if it had been found, would have important implications for our understanding of virus assembly. This is because the vast majority of *in vitro* studies with recombinant Gag protein have used a truncated protein lacking p6. The findings presented here thus provide important support for the relevance of these studies to the virus-assembly problem.

Multimerization of Gag upon nucleic acid binding has been previously investigated both *in vitro* and in cells (48, 49). However, the stronger tendency for Gag to multimerize on Psi RNA relative to non-Psi RNA has, to our knowledge, not been previously reported. The dimerization of Gag protein on Psi RNA would appear to have major ramifications regarding virus particle assembly.

Gag normally packages Psi-containing RNA with very high selectivity, despite the presence of a substantial excess of cellular mRNA species that can also be packaged. The mechanism of this selective packaging is not fully understood. We have previously reported (9, 10) that at physiological ionic strengths, Gag binds with similar, high affinities to RNAs with or without the packaging signal. Thus, the selection of Psi^+^ RNA during virus assembly is evidently not due to a uniquely high affinity of Gag for this RNA. We and others have suggested that binding to Psi induces a nucleating event, such as formation of a small Gag oligomer, more efficiently than binding to other RNAs, and that this difference might underlie the selective packaging of Psi^+^ RNA (11, 12, 50–52). The present results provide very strong support for this fundamental concept.

A speculative model for the nucleation of HIV-1 viral assembly, based on previous studies (10, 39, 53, 54) and the new work reported here, is shown in Figure 6. In this model, Gag exists in an equilibrium between bent and extended conformations. We previously hypothesized that different Gag conformations are adopted while binding to different RNAs (10). When Gag binds Psi, the extended conformation is favored, which allows rapid dimerization of Gag via CA-CA (38, 39), SP1-SP1 (55, 56), and potentially NC-NC interactions (49). In contrast, the bent conformation is preferred for binding to TARpolyA, requiring a rate-limiting conformational switch before dimerization can occur. In the bent conformation, MA binding to the nucleic acid may also further stabilize this non-productive complex. The capability of nMS to characterize disordered and heterogeneous systems has allowed us to gain new insights into viral assembly. In the future, additional MS-based technologies such as ion-mobility MS, RNA-protein crosslinking approaches, and covalent labelling may provide additional insight into RNA-induced Gag conformational changes and virion assembly. The versatility of nMS may also allow the future investigation of Gag binding to the complete 5′ UTR of gRNA.

**Figure 6.**
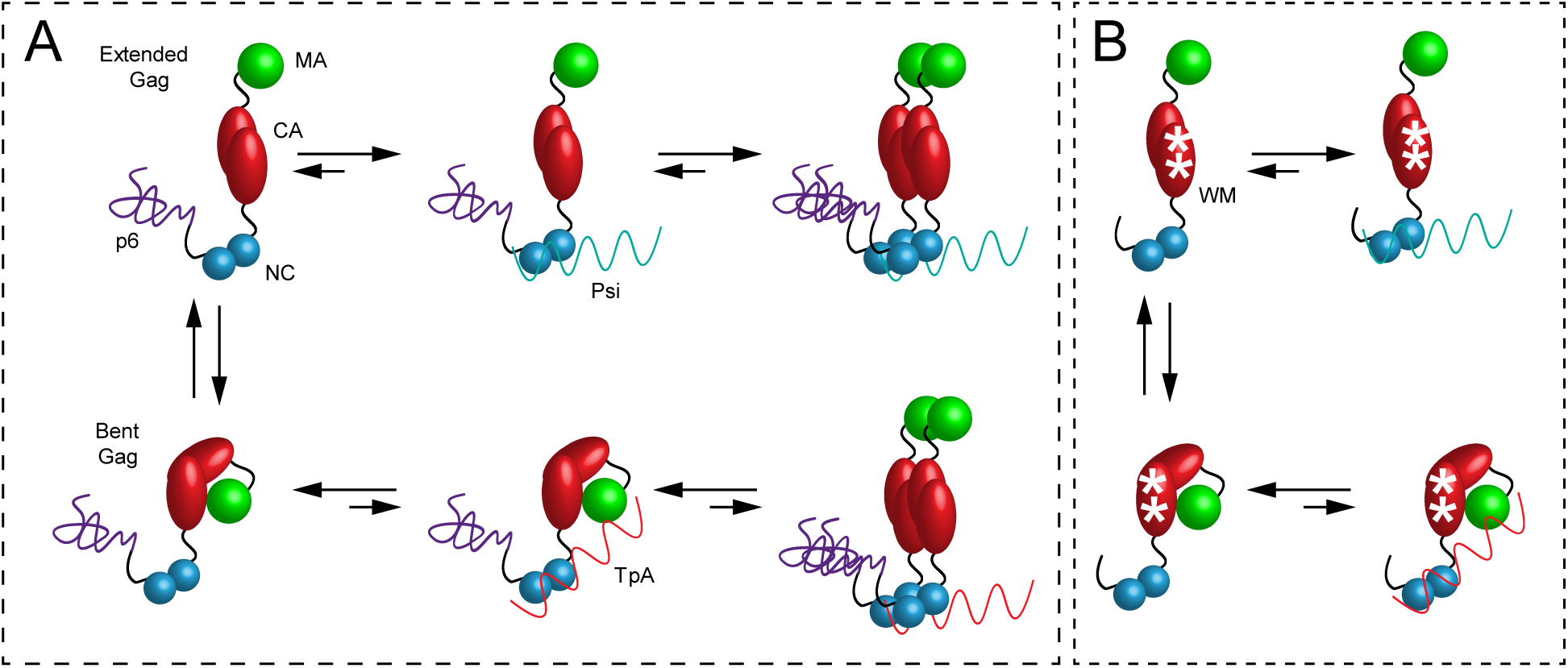
Model for WT (A) and mutant (B) WM-GagΔp6 binding to Psi RNA (top) and non-Psi RNA (*bottom*). In the absence of RNA, WT Gag is a monomer in equilibrium between bent and extended conformations. Gag binds Psi in a dimerization-competent extended conformation and dimerizes. Gag binds TpA in the dimerization-incompetent bent conformation and must undergo a rate-limiting extension before dimerization occurs.

## Experimental Procedures

### Protein purification

WT HIV-1 BH10 Gag, GagΔp6, and WM-GagΔp6 (Figure 1A) were expressed in *Escherichia coli* (BL21(DE3)pLysS) and purified as previously described (57) with the following alterations. Soluble lysate fractions were treated with polyethylenimine (PEI) to precipitate endogenous nucleic acids prior to the ammonium sulfate precipitation step. The pellet was resuspended in 100 mM NaCl, 20 mM Tris-HCl, pH 7.4, 10 mM 2-mercaptoethanol (βME) and 1 µM ZnCl_2_, loaded onto a HiTrap Heparin HP affinity column (GE Healthcare) equilibrated in the same buffer, and eluted with 0.75 – 1 M NaCl as described (58).

The purification of WT HIV-1 Gag was identical to that of GagΔp6 except for the addition of a cleavable C-terminal His_6_-tag. The Gag-His_6_ protein was loaded onto a HIS-select affinity column (Sigma), washed (20 mM Tris-HCl, pH 7.4, 500 mM NaCl, 5 mM βME, 1 µM ZnCl_2_, 0.1% Tween-20, 5 mM imidazole) and eluted in the same buffer using a step gradient of 10 mM, 20 mM, 50 mM, 75 mM and 150 mM imidazole. Gag eluted in the 20-75 mM imidazole fractions. These fractions were pooled and dialyzed overnight into the same buffer without imidazole using a Slide-A-Lyzer Dialysis Cassette (20k MWCO, Thermo Scientific). During dialysis, the His-tag was cleaved using 1 mg Tobacco Etch Virus (TEV) protease/20 mg of Gag. The TEV protease was purified in-house as described (59).

WT Gag was further purified by elution through a HiTrap Heparin HP affinity column as described above. All steps were carried out at 4 °C and in all cases, the purest protein fractions were pooled for use in salt-titration binding and nMS studies (Figure S10). For nMS studies, purified Gag proteins were dialyzed (3500 MWCO dialysis tubing, BioDesign Inc. of New York) into 500 mM ammonium acetate, pH 6.8 and concentrated using an Amicon-ultra 30k MWCO ultrafiltration device (Millipore-Sigma). For salt-titration binding assays, the proteins were dialyzed using Slide-A-Lyzer Dialysis Cassette (20k MWCO) into 20 mM Tris-HCl, pH 7.4, 500 mM NaCl, 10% glycerol, 10 mM βME, and 1 µM ZnCl_2_. Protein concentrations were determined by measuring the absorbance at 280 nm using an extinction coefficient of 63223 M^-1^cm^-1^.

### RNA preparation

HIV-1 NL3-4 TARPolyA and Psi RNAs shown in Figure 1B were generated via T7 RNA polymerase-directed *in vitro* transcription as described (60) from either a TARPolyA-encoding pUC19 plasmid or a Psi-encoding pIDTsmart plasmid (Integrated DNA Technologies). Plasmids were linearized via FOK1 digestion prior to transcription. TARPolyA was preceded by a hammerhead ribozyme designed to cleave during the transcription reaction, resulting in the desired 5′ end. A hammerhead ribozyme construct was only used in the case of TARpolyA as it significantly improved the homogeneity of this RNA, as analyzed by nMS. The Psi RNA was engineered with two additional guanosines at the 5’-end in order to improve the yield of *in vitro* transcription. This Psi construct was shown previously to be recognized by Gag with high specificity (10). RNAs were purified by electrophoresis on 6% polyacrylamide/8 M urea denaturing gels followed by elution and concentration via the crush and soak method (60). RNA quality was checked on 6% polyacrylamide/8M urea denaturing gels with ethidium bromide staining (not shown) and by MS (Figure S11).

### Fluorescence anisotropy binding assays

RNAs used in FA-based salt-titration binding assays were labeled with fluorescein-5-thiosemicarbazide (FTSC) at the 3’-end as described (42). The labeling efficiency of the RNAs was determined by measuring the absorbance at 260 nm and 495 nm and using the extinction coefficients ε_495nm_ = 8.5 × 10^4^ M^-1^·cm^-1^ for FTSC and ε_260nm_ = 9.3 × 10^5^ M^-1^·cm^-1^ and 9.7 × 10^5^ M^-1^·cm^-1^ for TARpolyA and Psi RNAs, respectively. A correction factor (ε_260nm_/ε_495nm_ = 0.3266) was used to correct the RNA concentration for the absorbance of the dye at 260 nm. The NaCl salt-titration binding assays were performed and the data analyzed as described (42). A very similar approach was used to determine if Gag:RNA binding was affected by nMS-compatible solution conditions. For these measurements, all non-volatile salts were replaced with ammonium acetate in both the RNA and the Gag samples. FTSC-labeled RNAs were refolded in 50 mM ammonium acetate, pH 6.8 and 1 mM magnesium acetate by heating (90 °C, 2 min) and snap-cooling (ice, 2 min) followed by incubation at 37 °C for 30 min. The GagΔp6 protein was dialyzed overnight in 500 mM ammonium acetate, pH 6.8, prior to carrying out the ammonium acetate salt-titration binding assays and data analysis as previously described (42).

### Native mass spectrometry

Analysis of the three Gag constructs (WT Gag, GagΔp6, and WM-GagΔp6) in the absence of RNA was carried out at both 3 and 16 µM protein in 500 mM ammonium acetate, pH 6.8. Prior to nMS analysis, RNAs were refolded as described above. Gag:RNA complexes were formed by mixing 3 µM Gag with 0.5 µM refolded RNA at room temperature. Gag samples were stored on ice prior to analysis while reactions containing Gag and RNA were incubated at room temperature for 15-30 minutes prior to analysis. An aliquot (1-3 µL) was loaded into a borosilicate glass nano-electrospray ionization (nESI) emitter prepared in house on a Sutter p-97 micropipette puller and directly injected into a ThermoFisher Scientific Q Exactive(tm) UHMR Hybrid Quadrupole-Orbitrap(tm) mass spectrometer. Separate nESI emitters were used for each measurement. The instrument settings were as follows: spray voltage, 0.9-1.1 kV; capillary temperature, 225 °C; S-lens RF level, 200; High energy collision dissociation (HCD), 0 V; in-source fragmentation, 0 V; in-source trapping, -100 V; trap gas, 8 (arbitrary units); m/z range, 500-80,000. A minimum of three technical repeats were performed for each combination of Gag and RNA. Mass spectra were deconvolved using UniDec v4.1 (61, 62). Theoretical molecular weights were calculated based on protein and RNA sequences (provided in Table S2) using the Protein/RNA molecular weight calculator tool in UniDec.

### Constructs for analysis of RNA packaging

Gag was expressed in mammalian cells from pCMV55M1-10 (63), a Rev-independent version of the HXB2 Gag gene. This plasmid does not contain Psi. We measured packaging of the RNA from an HIV-1-derived GFP vector constructed from pLenti6/V5-DEST (ThermoFisher, Inc.). A Psi-version of this vector was constructed, using inverse PCR, by deleting nucleotides corresponding to 214-365 of NL4-3 RNA; the deleted stretch encompasses the stem-loops SL1-SL4. Mutants of the Gag gene in pCMV55M1-10 were generated by inverse PCR. All constructs were confirmed by sequencing of the entire coding region.

### Transfection

HEK293T/17 cells were seeded at a density of 4 × 10^6^ cells in a 10 cm cell culture dish. The following day they were transfected with 6 μg Gag plasmid + 6 μg vector plasmid + 3 μg pCMV-Rev (to support nuclear export of vector RNA) using Transit-293 (Mirus) following the manufacturer’s protocol. Cell culture supernatants were collected after 48 hr and 72 hr and filtered through 0.22 µm syringe filters (Merck Millipore). Filtered supernatants were stored at -80 °C until further analysis. The transfected cells were lysed at 72 hr for immunoblotting as described below. The results presented represent three independent transfections.

Transfected cultures were also analyzed by transmission electron microscopy. Cells were fixed 48 hr after transfection and processed as described previously (55).

### RNA analysis

For extracting RNA from VLPs, the filtered cell culture fluid was ultracentrifuged (25k rpm, 4 °C, SW55Ti Rotor, Beckman Coulter) through a 20% sucrose cushion prepared in TNE buffer (10 mM Tris-HCl, 100 mM NaCl and 1 mM EDTA, pH 7.5). The VLPs in the pellet were lysed by adding PK Lysis buffer (50 mM Tris-HCl, 100 mM NaCl, 10 mM EDTA, 1% SDS, 100 µg/ml Proteinase K, pH 7.5) and incubating at 37 °C for 30 mins. RNA was extracted from the lysed VLPs using TriReagent (Ambion) following the manufacturer’s protocol. Glycoblue (Ambion) was used as a carrier in the precipitation step. The RNA pellet was resuspended in nuclease-free water and stored at -80 °C until further analysis.

Copies of sequences in the RNA preparations were then enumerated by real-time RT-PCR. RNA representing ∼ 350 μl of culture fluid was treated with DNaseI (RNase free, Ambion) at 37 °C for 30 min in a total reaction volume of 10 μl. The DNaseI was inactivated by adding 1μl of 50 mM EDTA to the reaction followed by heat inactivation at 75 °C for 10 mins. First strand synthesis was performed using an iScript cDNA synthesis kit (Bio-Rad) in a total reaction volume of 20 μl following the manufacturer’s protocol. This DNA was then analyzed by SYBR Green-based (FastStart Essential DNA Green Master, Roche) real-time PCR. Standard curves were generated from serial dilutions of a transcript containing the target sequence. These transcripts were digested with DNaseI and the transcripts were cleaned up twice using RNeasy spin columns (Qiagen) after *in vitro* transcription. RNA transcripts were quantitated using A_260_ and Ribogreen (ThermoFisher, Inc.). In the real-time RT-PCR assays, HIV-1-derived packageable RNA was amplified using primers: 556 F: 5’-AAACAAAAGTAAGACCACCGCAC-3’ and 556 R : 5’-ACCACTCTTCTCTTTGCCTTGG-3’, spanning nucleotides 891-1047 of pLenti6/V5-DEST RNA. All real-time RT-PCR assays also included a no-RT control, which gave extremely low RNA copy numbers, indicating that the experimental values represent RNA copy numbers without significant DNA contamination.

### Immunoblotting

Gag in pelletable VLPs was quantitated as follows. VLPs were pelleted through a 20% sucrose cushion in TNE buffer. Pellets were resuspended in 1XNuPAGE sample dye containing reducing agent and 1X HALT protease inhibitors (Invitrogen) and stored at -80 °C until further analysis. For detecting intracellular HIV-1 Gag, cells were lysed using the same 1X NuPAGE sample dye cocktail (Invitrogen), sonicated for complete lysis and stored at -80 °C. Prior to electrophoresis samples were thawed and heated at 90 °C for 5 mins. Electrophoresis was carried out using NuPAGE 4 to 12% Bis -Tris polyacrylamide gels (Invitrogen) in 1X NuPAGE buffer followed by transfer to Immobilon-FL polyvinylidene difluoride membrane (Millipore). Membrane was blocked using Intercept Blocking Buffer (LI-COR); this was followed by the addition of primary antibodies (goat anti-p24 from NIH, and mouse anti-actin, Santa Cruz Biotechnology, C4 sc-47778) diluted in the blocking buffer and incubated overnight at 4 °C. Membranes were washed thrice with TBS (20mM Tris-HCl, pH 7.5/500 mM NaCl) and probed with secondary antibodies conjugated to Dylight 800 or 700 (LI-COR). Blots were imaged using the Li-COR Odyssey imaging system and images were analyzed using ImageStudioLite. Absolute quantities of Gag protein were obtained by reference to a standard curve prepared using recombinant GagΔp6 (a kind gift of S. Datta, NCI), an example of which is in Figure S12.

Sample band intensities were within the linear range of the standard curve.

## Data Availability

All data are contained within the article.

## Acknowledgements

We thank Drs. Zac VanAernum and Florian Busch for mass spectrometry assistance and discussion and Drs. Ioulia Rouzina and Siddhartha Datta for helpful discussions. We also thank Ferri Soheilian and Kunio Nagashima for electron microscopy and Dr. Barbara Felber (NCI-Frederick) for the gift of the pCMV55M1-10 and pCMV-Rev plasmids.

